# PredIDR: Accurate prediction of protein intrinsic disorder regions using deep convolutional neural network

**DOI:** 10.1101/2024.07.24.604908

**Authors:** Kun-Sop Han, Su-Jong Yun, Chol-Song Kim, Chol-Pyok Ri, Alessio Del Conte, Damiano Piovesan

**Author notes:** Correspondence: Damiano Piovesan, Department of Biomedical Sciences, University of Padova, Padova, Italy.

## Abstract

The involvement of protein intrinsic disorder in essential biological processes, it is well known in structural biology. However, experimental methods for detecting intrinsic structural disorder and directly measuring highly dynamic behavior of protein structure are limited. To address this issue, several computational methods to predict intrinsic disorder from protein sequences were developed and their performance is evaluated by the Critical Assessment of protein Intrinsic Disorder (CAID). In this paper, we describe a new computational method, PredIDR, which provides accurate prediction of intrinsically disordered regions in proteins, mimicking experimental X-ray missing residues. Indeed, missing residues in Protein Data Bank (PDB) were used as positive examples to train a deep convolutional neural network which produces two types of output for short and long regions. PredIDR took part in the second round of CAID and was as accurate as the top state-of-the-art IDR prediction methods. PredIDR can be freely used through the CAID Prediction Portal available at https://caid.idpcentral.org/portal or downloaded as a Singularity container from https://biocomputingup.it/shared/caid-predictors/.

## 1. Introduction

Intrinsically disordered regions (IDRs) in proteins lack a fixed three-dimensional structure under physiological conditions and perform their functions without fully folding^[10,16,24]^. According to recent studies, IDRs are abundant in living things and are found in more than 30% of proteins, particularly in the eukaryotes^[25, 29]^. Moreover, proteins with IDRs are involved in a variety of important biological processes^[28]^.

Structural biology experimental methods, including X-ray crystallography, circular dichroism and nuclear magnetic resonance spectroscopy, are used for deriving information about intrinsic disorder. However direct measurements of such highly dynamic behavior is still difficult^[5]^. Since only a few thousands of IDRs were experimentally characterized^[12]^, numerous computational methods to predict IDRs from protein sequences were developed over the years^[6,14,17,21]^. These methods have been proven useful for structural and functional characterization of specific proteins^[8,18]^ and for analyses at proteome scale^[9,13,26,27,30]^.

Performance of disorder predictors was assessed in community-driven competitions including the Critical Assessment of protein Structure Prediction (CASP) and more recently the Critical Assessment of protein Intrinsic Disorder (CAID). During the six experiments from CASP5 to CASP10, where disorder was evaluated considering X-ray missing residues, the number of disorder predictors increased from 6 ^[20]^ to 28^[23]^. Following CASP10, the assessment of the disorder was provided by CAID organizers. In CAID, both the ability of predicting IDRs and the binding sites within IDRs, are evaluated. CAID participants submit their prediction software to the organizers, who generate predictions by running the program on the given protein targets whose IDR (and binding site) annotations were not available at the training time. In CAID, the task of a disorder (or binding site) predictor is to provide a score to each amino acid residue representing the probability to be intrinsically disordered or to be in a binding site. In the first two CAID editions, which were organized in 2018 and 2022, 43 and 71 prediction methods were evaluated^[4,5]^, showing an increasing attention to the CAID challenge.

Recently, the CAID Prediction Portal, a web server which executes all CAID methods, was developed^[1]^. This comprehensive service generates a standardized output, and provides the users with the convenient environment to facilitate comparing methods and produce a consensus prediction.

According to the latest CAID2 assessment^[5]^, the performances of different IDR prediction methods vary across different benchmarks, which emphasizes the need for continued development of more versatile and efficient prediction softwares.

In this work, we present a new deep convolutional neural network, PredIDR, which accurately predicts intrinsically disordered regions in proteins, reflecting and capturing the features of X-ray missing residues as available in Protein Data Bank (PDB) structures^[3]^. In the following, firstly we examine the effect of various feature combinations. Next, we assess the impact of our ensemble approach and smoothing techniques. Finally, we compare our methods with the state-of-the-art IDR prediction methods participating in CAID2 on different testing sets.

## 2. Materials and Methods

### 2.1. IDR analysis of non-redundant sequences extracted from PDB

We obtain 8,657 non-redundant, high-resolution protein sequences from the PDB database (August 09, 2019). Proteins were filtered so that their sequence identity is lower than 25% (using CD-HIT) and they are longer than 51 residues (see Additional file 1). In our experiment, a disordered residue is inferred from the missing residues in X-ray experiments. That is, a disordered residue is defined as the residue without 3-dimensional (3D) coordinates despite being incorporated in the crystal. We considered only those IDR segments of at least four consecutive disordered residues. Of the 8,657 non-redundant PDB chains, 5,997 (called “IDR chain”, see Additional file 2) have at least one IDR and 2,660 (called “STR chain”, see Additional file 3) are fully structured.

### 2.2 Validation, testing and training set

#### 2.2.1 Validation set

We randomly divided 5,997 IDR chains into 2 datasets: one (called Train5400, see Additional file 4) includes 5,400 chains, which will be used to make a training set together with 2,660 STR chains, and the other includes 597 chains, which constitute a validation dataset (called Val597, see Additional file 5).

Val597 dataset contains 151,203 residues and 950 disordered regions in 597 chains (Table 1). Of 151,203 residues, 12,014 (7.95%) residues are positives, 139,189 (92.05%) residues are negatives.

**Table 1.**
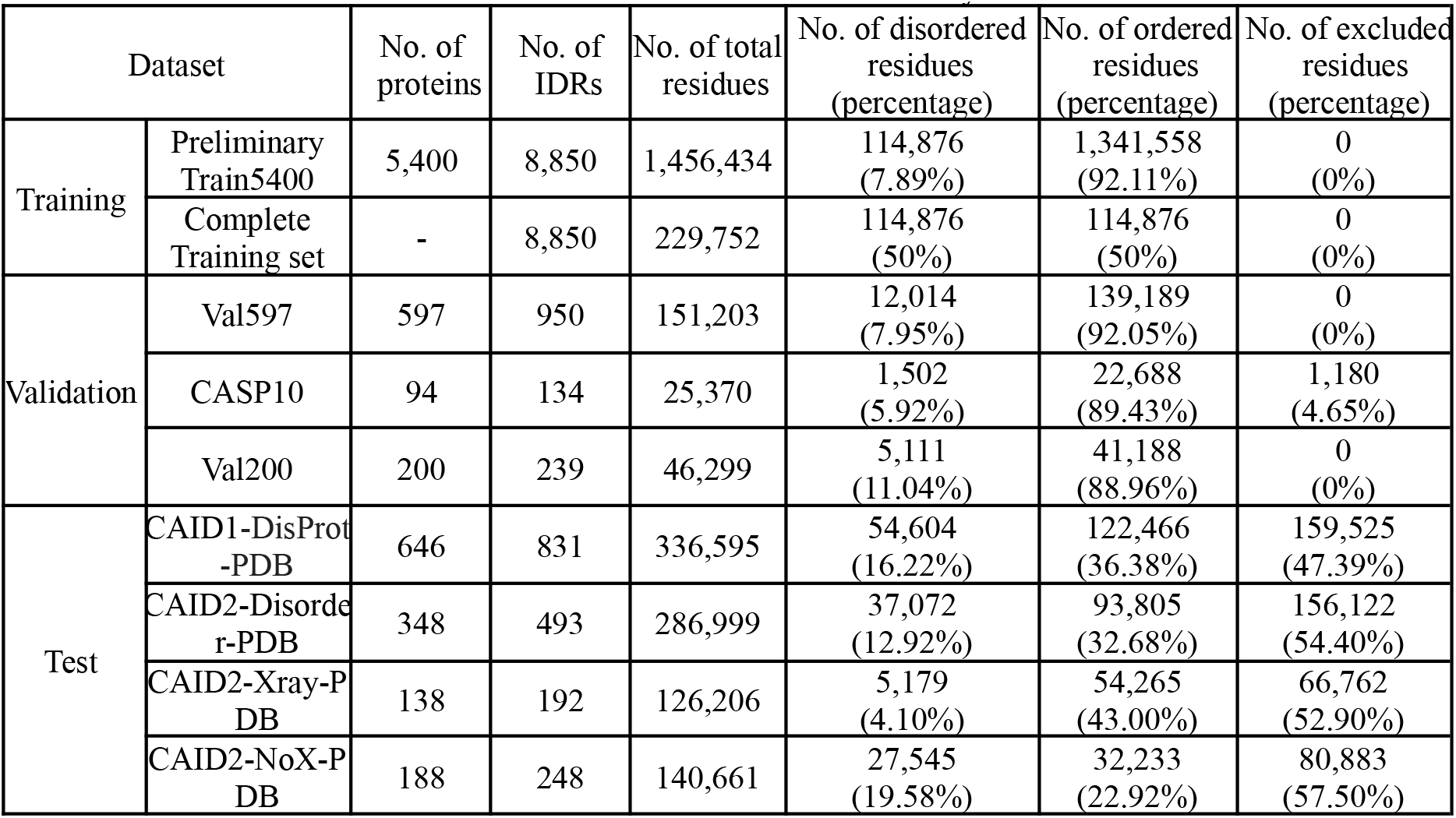
Datasets used in this study.

In addition to Val597 dataset, we use another two validation sets (Table 1): one is CASP10 dataset (see Additional file 6) which was released in IDR category of CASP10^[23]^ and the other is Val200 set (see Additional file 7), which consists of 200 protein chains with only the IDRs of at least ten residues, extracted from Val597 dataset. Val200 was created to mimic the CAID datasets, whose targets contain only the IDRs of at least ten residues. CASP10 dataset contains 25,370 residues and 134 disordered regions in 94 chains. Of 25,370 residues, 1,502 (5.92%) are positives, 22,688 (89.43%) are negatives and 1,180 (4.65%) are excluded for evaluation as per CASP10 assessors decision. Val200 dataset contains 46,299 residues and 239 disordered regions in 200 chains. Of 46,299 residues, 5,111 (11.04%) are positives, 41,188 (88.96%) are negatives.

#### 2.2.2 Testing set

Before CAID2 experiment, we used DisProt-PDB dataset (downloadable at https://caid.idpcentral.org/challenge/results) of CAID1 as a testing set, which was released in CAID1 experiment^[4]^. To avoid confusion with the Disorder-PDB dataset of CAID2, we have called it CAID1-DisProt-PDB (see Additional file 8) in this paper. The disordered residues are positives, the residues which are observed in PDB and which are not disordered are the negatives, and the remaining residues are excluded in all CAID-related testing sets. The CAID1-DisProt-PDB dataset contains 646 sequences, 336,595 residues and 831 disordered regions (Table 1). 54,604 (16.22%) residues are contained in 831 disordered regions, 122,466 (36.38%) residues are labeled as negatives and 159,525 (47.39%) are excluded.

During CAID2 experiment, our prediction methods were evaluated on the Disorder-PDB dataset (downloadable at https://caid.idpcentral.org/challenge/results) of CAID2^[5]^, which is called here CAID2-Disorder-PDB (see Additional file 9). The dataset contains 286,999 residues and 493 disordered regions in 348 sequences (Table 1). Of 286,999 residues, 37,072 (12.92%) are positives, 93,805 (32.68%) are negatives and 156,122 (54.40%) are excluded.

After publication^[5]^ of the CAID2 result, we created two additional test sets, CAID2-Xray-PDB and CAID2-NoX-PDB (Figure 1) to inspect the predictor ability to detect two different flavors of disorder.

**Figure 1.**
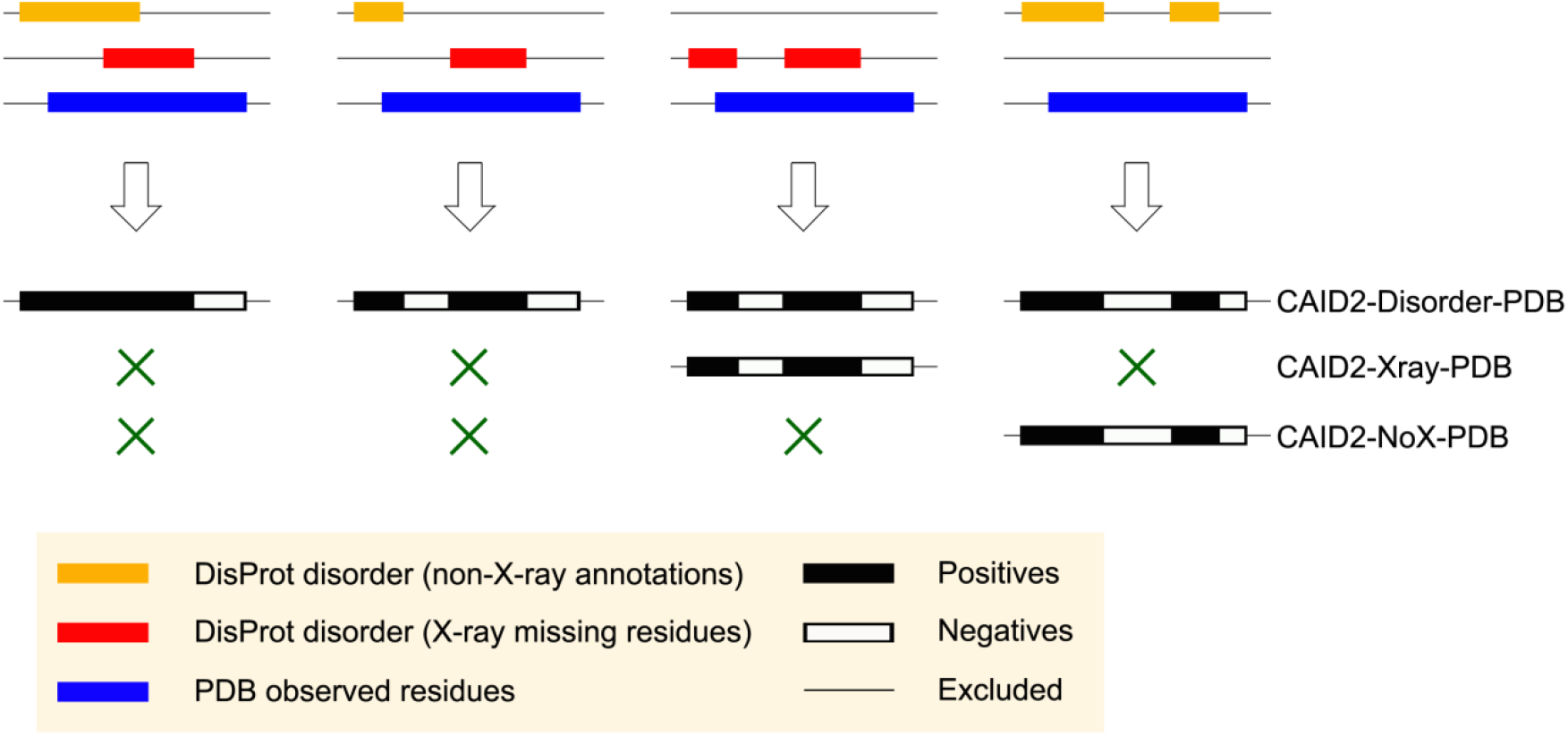
The three CAID2 test sets. Different data sources are combined to generate the CAID2-Disorder-PDB, CAID2-Xray-PDB and CAID2-NoX-PDB test sets. Red and orange boxes represent disordered regions annotated by X-ray experiment and all disorder annotations except X-ray missing residues, respectively.

Figure 1 shows four possible ways of combining different sources of data, two for disorder (orange, red) and one for structure (blue). The first case shows the overlap of disordered regions annotated by two origins (X-ray crystallography and all disorder annotations except X-ray missing residues (called non-X-ray annotations)). The second case indicates that the two different kinds of disordered regions are away from each other. The third case contains only disordered regions annotated by X-ray missing residues. The fourth case shows disordered regions annotated only by non-X-ray annotations.

The CAID2-Disorder-PDB dataset released in the CAID2 experiment is constructed from all four possible cases of Figure 1.

In order to test the ability of disorder predictors to detect X-ray missing residues from protein sequences, we built “CAID2-Xray-PDB” dataset (see Additional file 10) containing only X-ray missing residues as disorder. This dataset is generated by subtracting the Disorder-NOX^[5]^ dataset (all disorder annotations except Xray missing residues) to the CAID2-Disorder-PDB. The CAID2-Xray-PDB dataset contains 126,206 residues and 192 disordered regions in 138 sequences (Table 1). Of 126,206 residues, 5,179 (4.10%) are positives, 54,265 (43.00%) are negatives, and 66,762 (52.90%) are excluded.

We also want to examine the ability of disorder predictors to detect the non-X-ray annotations-based disorder regions from protein sequences. For this purpose, we made another dataset called “CAID2-NoX-PDB” (see Additional file 11). Similarly to the CAID2-Xray-PDB dataset, it is produced by comparing the non-X-ray annotations-based disorder regions of Disorder-NOX^[5]^ dataset and the disorder regions of CAID2-Disorder-PDB dataset and collecting the sequences containing only the non-X-ray annotations-based disorder residues as disorder. This dataset contains 140,661 residues and 248 disordered regions in 188 sequences (see Table 1). Of 140,661 residues, 27,545 (19.58%) are positives, 32,233 (22.92%) are negatives, and 80,883 (57.50%) are excluded.

#### 2.2.3 Training set

Predicting whether a protein residue is disordered or ordered is a binary classification problem. For such problems, some machine learning techniques such as shallow neural networks, support vector machines and deep neural networks can be employed, where a better regression is expected when the number of positive labels is equal or similar to the number of negative ones in the training set.

However, as shown in Table 1, the Train5400 dataset is very unbalanced with only 7.89% of disordered residues.

In order to provide the same number of positive and negative examples, we built an artificial dataset where negative examples are undersampled in order to obtain a perfectly balanced dataset.

Table 2 shows the distribution of disordered and structured (non-IDR) regions in the sequence. As shown in Figure 2A, the IDR_N subset (see Additional file 12) includes IDRs starting at N-terminus of the protein chain and ending in the middle while IDR_C are IDRs starting in the middle and ending at C-terminus of the protein chain (see Additional file 13). The IDR_M subset (see Additional file 14) includes IDRs which are neither starting at the N-terminus nor ending at the C-terminus of the protein chain, i.e. are fragments in the middle of the protein. All 8,850 IDRs belong to the three IDR subsets, because there are no fully disordered proteins in the Train5400 dataset.

**Table 2.**
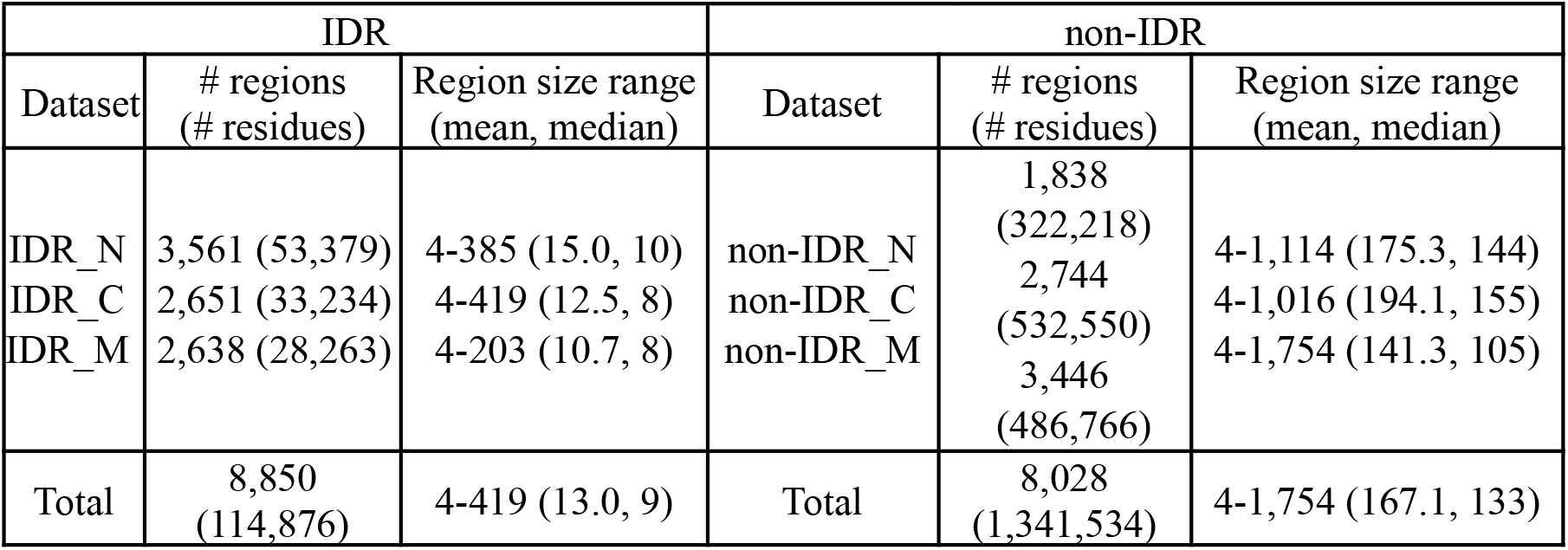
Distribution of IDRs and non-IDRs sequence position in the Train5400 dataset.

**Figure 2.**
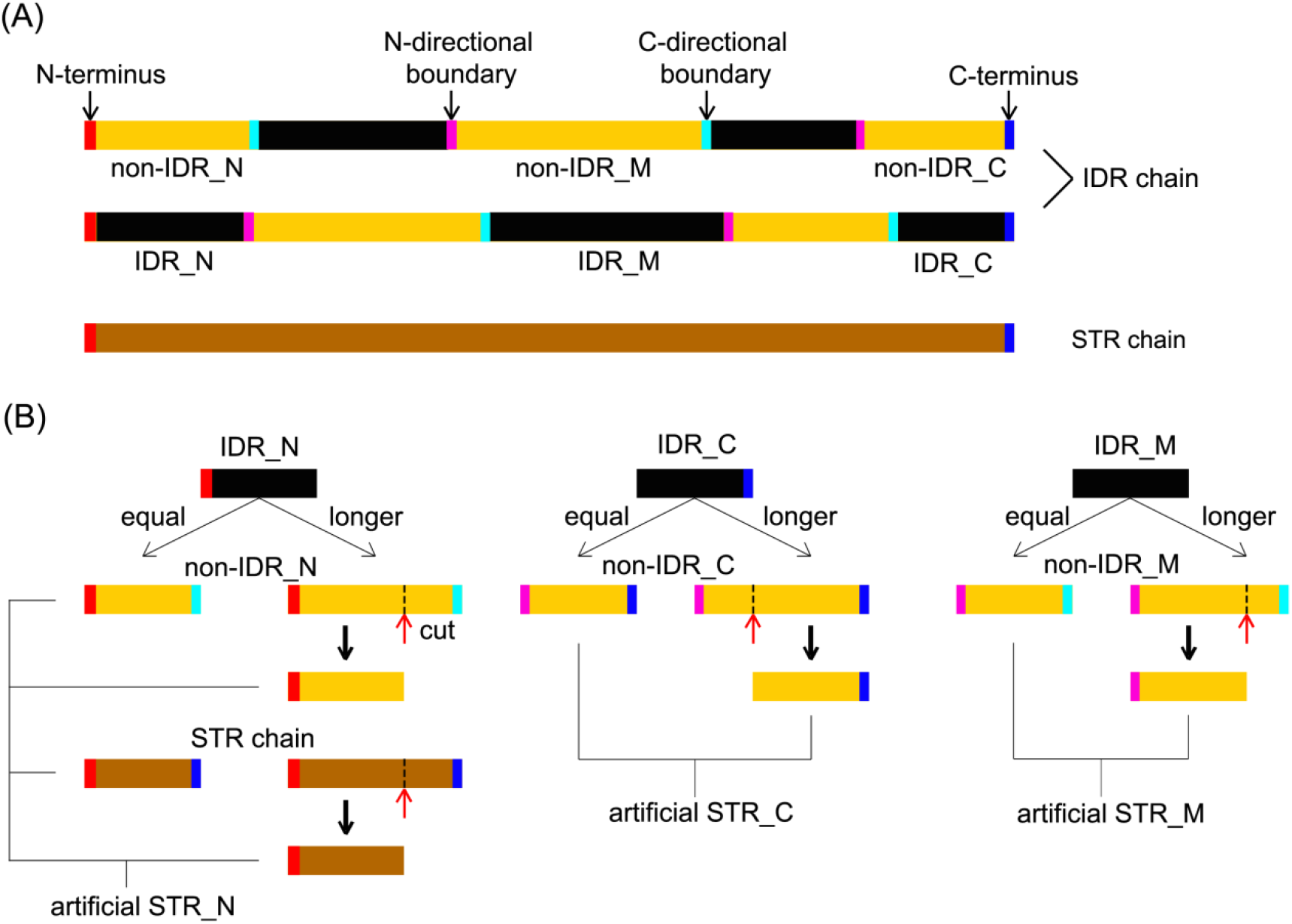
The process of making artificial STR_N, STR_M and STR_C. (A): Two IDR chains and a STR chain. IDR_N, IDR_C and IDR_M segments are coloured black and non-IDR_N, non-IDR_M and non-IDR_C segments are coloured orange. N-terminus and C-terminus are coloured red and blue, respectively. N- and C-directional boundaries are coloured pink and light blue, respectively. An IDR chain has any of three kinds of IDRs. A STR chain coloured brown has no any IDRs. (B): The process of making artificial STR_N, STR_M and STR_C. Artificial STR_M and STR_C can be made from non-IDR_M and non-IDR_C, respectively. C-terminus is preserved when making artificial STR_C from non-IDR_C. Whether N- or C-directional boundary is preserved is randomly chosen when making artificial STR_M from non-IDR_M. An artificial STR_N can be made from a non-IDR_N or a STR chain. N-terminus is preserved when making the artificial STR_N. The red arrow indicates the cutting position.

As shown in Table 2, there are 8,028 non-IDRs (constitute a non-IDR set) in the Train5400 dataset. Similar to the IDR set, the non-IDR set is divided into three subsets according to their position in the protein sequence: non-IDR_N, non-IDR_C and non-IDR_M subset (see Additional file 15, 16 and 17 and Figure 2A). Non-IDRs with less than four residues were excluded. All 8,028 non-IDRs belong to the three subsets as all protein sequences of the Train5400 dataset contain at least one IDR.

In order to balance the number of regions and residues in the training dataset for the three categories described, we applied the following algorithm (Figure 2B and Figure 3).

**Figure 3.**
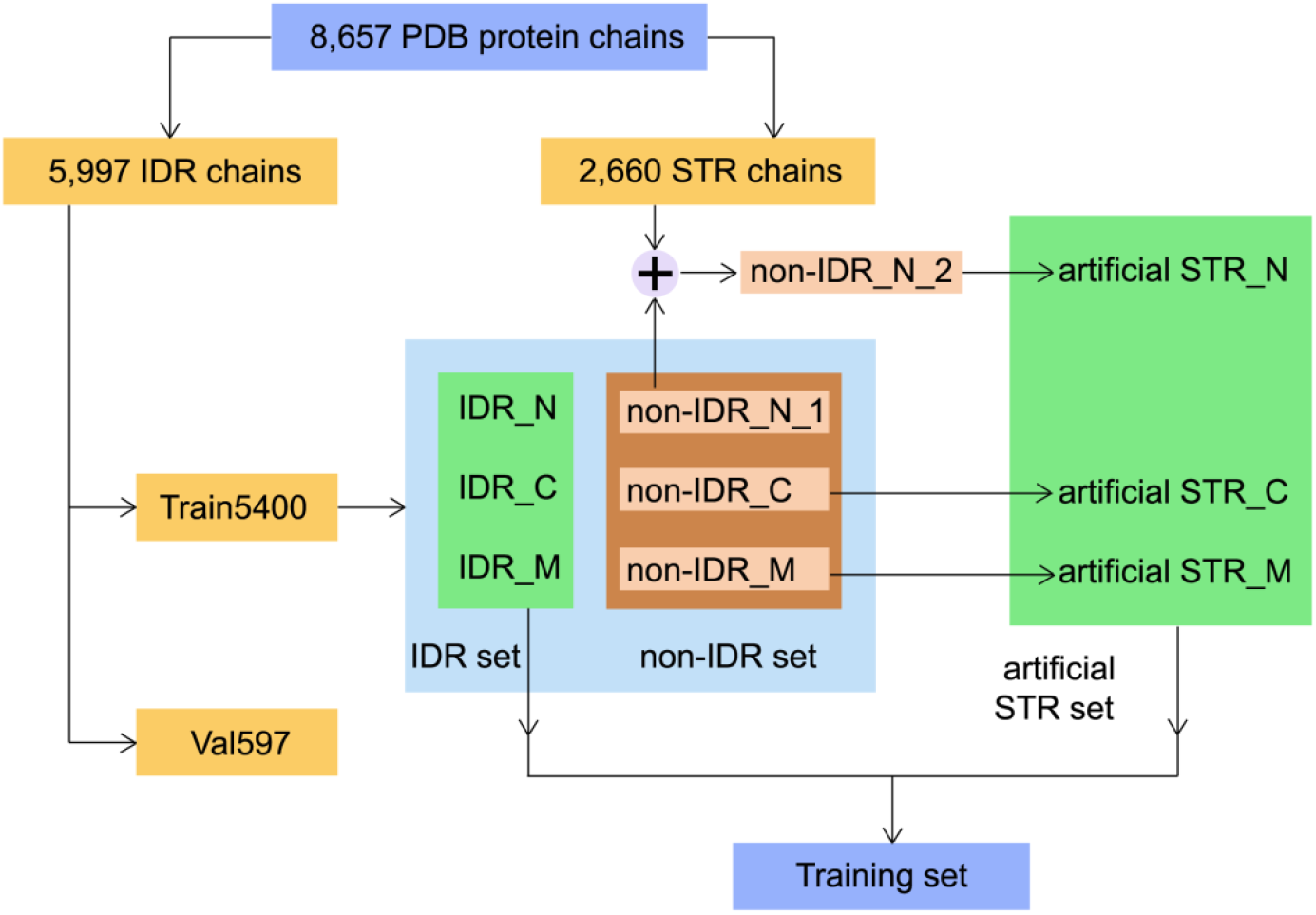
An overall architecture of constructing the Training set from 8,657 protein chains.

For non-IDR_C and non-IDR_M subsets we simply sampled available regions in each category in order to obtain exactly the same length distribution in the corresponding positive IDR_C and IDR_M classes (Figure 3). The sampling was performed first matching regions with the same length and then sorting by length both positive and negative remaining regions, pairing the two lists and selecting the closests in length. When structured regions (negatives) are longer than the paired disordered regions (positives) they are cut to match the length of the disordered counterpart (Figure 2B). C-terminus is preserved when making an artificial STR_C from a non-IDR_C. In the case of non-IDR_M, the decision about cutting towards the N- or C-termini is random. For the non-IDR_N category, however, the number of non-IDR_N regions (1,838, named non-IDR_N_1) is smaller than IDR_N regions (3,561) (Table 2). Therefore, we created additional artificial non-IDR_N regions drawing from STR chains (Figure 2B and Figure 3). N-terminus is preserved when making an artificial STR_N from a non-IDR_N or a STR chain (Figure 2B).

Finally, our training set consists of 114,876 positives (114,876 IDR residues from 8,850 IDRs) plus 114,876 negatives (114,876 STR residues from 8,850 artificial STRs) (Table 1).

### 2.3 Input features

In our experiments, we represent each protein residue with three different features, evolutionary profile, secondary structure and solvent accessibility. Sequence profiles are obtained from a multiple sequence alignment (MSA) generated by a PSI-BLAST^[2]^ search (e=0.001, 3 iterations) against UniRef50 sequence database (as available in SCRATCH 1.2^[19]^), and retaining only those MSA sequences with a coverage of more than 50% with the target sequence. Secondary structure is obtained by SCRATCH 1.2, which produces both three-state (SS3) and eight-state (SS8) secondary structure predictions. Solvent accessibility is also predicted by SCRATCH 1.2, which produces both binary (buried/exposed, ACCBIN) and 20-class (ACCDEC) predictions.

### 2.4 Encoding scheme

Since the disorder status of a residue can be influenced by neighboring residues, we used a sliding window. All features in the sliding window are combined to form a feature tensor. The size of the tensor for each position in the window is 32 consisting of 21 frequency values (one for each amino acid type plus the gap symbol) as provided by the MSA profile, 8-dimensional vector for the 8-state secondary structure (SS8), one value for the predicted solvent accessibility (ACCDEC) and 2 values indicating whether the position is outside the sequence boundaries, i.e. before N-terminus or after C-terminus. In our experiments, we used a window of 15 residues, therefore, the size of the feature tensor for a single residue is 32 by 15.

We used only one output node to predict the state of a residue where the output is 1 if the residue is disordered and 0 if predicted as ordered.

### 2.5 Architecture of 2D convolutional neural network employed in PredIDR

We score a sliding window centered on a given residue using a 2-dimensional (2D) convolutional neural network (Figure 4). The architecture of the network consists of an input layer, two 2D convolutional layers with rectified linear unit (ReLU) activation, a 2D max-pooling layer, one fully connected layer with ReLU activations and one fully connected layer with no activations.

**Figure 4.**
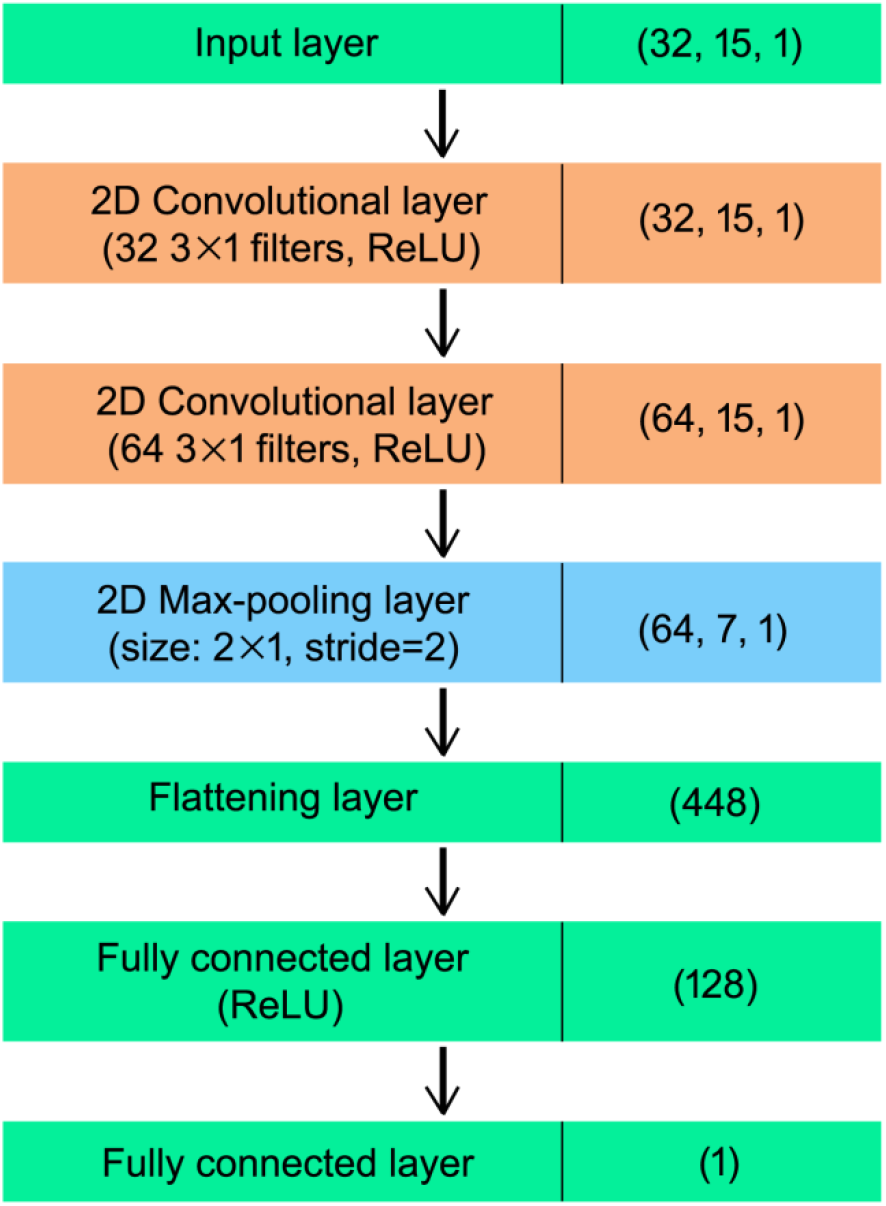
The architecture of the 2D convolutional neural network used in PredIDR. The values in brackets of the right column indicate the dimensionality of the output of each layer. The order of the three values is: (*number of channels, height, width*).

The input layer has size (32, 15, 1) corresponding to the size of the sliding window and the number of encoded features and the output of the network is a single node generating a number representing the structural propensity of the central residue of the window to be disordered..

The input map is processed by a 2D convolutional layer with 32 filters of size 3×1 which produces 32 output feature maps of size 15×1. 3×1 filters are used in two 2D convolutional layers as the height and width of the input feature are 15 and 1 respectively. A second 2D convolutional layer with 64 filters of size 3×1 generates 64 output feature maps of size 15×1. Rectified linear unit (ReLU) activation is used in two 2D convolutional layers. Then, these output feature maps are max-pooled with 2×1 windows and stride 2 such that 64 output feature maps with size 7×1 are produced. The flattening layer reshapes the feature maps of size 64×7×1 into a single vector of size 448. Afterwards, we compute several transformations using two fully-connected layers to produce the final output of the network. ReLU activation is provided in the first fully-connected layer with 128 neurons but not in the last fully-connected layer with only one neuron.

### 2.6 Training of convolutional neural network model

We trained our convolutional neural network on 72 examples per weight update, i.e. minibatch size is 72. As the training set contains 229,752 examples, 3,191 mini batches have been produced.

The parameters of the network model are trained using the Adam algorithm^[15]^. The Adam algorithm updates the parameters of the network by minimizing the mean squared error (mse) loss function (eq. 1).

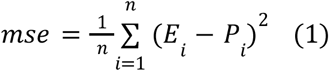

where *E*_*i*_ is the expected output for a given residue i (presence/absence of disorder represented as 1/0), *P*_*i*_ ∈ [0, 1] is the predicted output for the residue (representing the probability of a residue to be disordered) and n is the number of examples in a mini batch or a set of interest.

The initial learning rate for Adam was 0.005. The learning rate is divided by 10 after every 10 epochs. We typically get the best performance in 15–30 epochs, with the longest optimization epoch being 40.

### 2.7 Ensemble and smoothing

The generalization errors of neural network models can be minimized by using an ensemble predictor^[11]^. In our experiment, we trained the convolutional neural network twenty times and the ten top-performing networks were chosen based on the performance in Val597 dataset and combined as an ensemble by averaging the predictions from ten top-performing neural networks.

Finally, we smoothed the outputs of the ensemble by averaging the predictions of a sliding window of size w centered on the interested residue. We tested the effect of the smoothing in window sizes ranging from 3 to 23 for three validation sets.

### 2.8 Evaluation criteria

We used two probability-based evaluation measures (AUC_ROC and AUC_PR) which were used in the CASP10 assessment^[22]^. We briefly discuss those measures in the following.

We performed the receiver operating characteristic (ROC) and the precision-recall (PR) curve analysis for assessment of predictors’ ability to assign per-residue disorder confidence scores. For a set of thresholds (0 to 1), if the predicted probability of a residue is equal to or bigger than the threshold, it is considered as a disordered residue. A plot called a ROC curve is drawn between the true positive rates and false positive rates of a predictor over all thresholds. The area under the ROC curve (AUC_ROC) is used as a measure of the overall performance of a disorder predictor. A value of 1 indicates a perfect predictor, while 0.5 corresponds to a random prediction.

The PR curve is conceptually similar to the ROC curve^[7]^, but differs in that it is plotted between recall (TP/(TP + FN)) and precision (TP/(TP + FP)) coordinates. Similar to ROC curve analysis, the area under the PR curve (AUC_PR) is another probability-based measure of the disorder predictor’s accuracy. A value of 1 indicates a perfect performance of the disorder predictor.

## 3. Results and Discussion

We developed a new computational method, PredIDR, which produces the accurate prediction of the intrinsically disordered regions in proteins, in particular, the X-ray missing residues. In our method, the missing residues in X-ray crystallography in Protein Data Bank (PDB) were used as positive examples and we implemented a deep convolutional neural network which provides an estimate of the propensity of a protein residue to be disordered. In the following, we examined the effect of various feature combinations on the performance of PredIDR and assessed the impact of ensemble and smoothing procedures. Finally, we evaluated the performance of PredIDR on different testing sets.

### 3.1 Performance of different feature combinations

We used three different features including the sequence profile, the predicted secondary structure and solvent accessibility in our deep convolutional neural network. Table 3 shows the performance of various feature combinations on the Val597 dataset.

**Table 3.** Performance of different feature combinations on the Val597 dataset.

| Feature combinations | AUC ROC | AUC PR |
| --- | --- | --- |
| Profile | 0.841 | 0.462 |
| SS3 | 0.835 | 0.389 |
| SS8 | 0.875 | 0.557 |
| ACCBIN | 0.778 | 0.354 |
| ACCDEC | 0.921 | 0.604 |
| ACCDEC + SS8 | <u>0.939</u> | <u>0.703</u> |
| ACCDEC + profile | 0.928 | 0.698 |
| ACCDEC + SS8 + profile | **0.940** | **0.729** |

ACCDEC has AUC_ROC of 0.921 and AUC_PR of 0.604, which are the best performance of all the five single features. The 8-state secondary structure (SS8) performs better than the 3-state secondary structure (SS3) and the 20-class solvent accessibility prediction (ACCDEC) was more effective than the predicted binary solvent accessibility (ACCBIN). To avoid redundancy, we chose the 8-state secondary structure and 20-class solvent accessibility.

Combining ACCDEC with any of the SS8 and profile features improved prediction performance than using ACCDEC alone.

The combination of three features (ACCDEC, SS8 and profile) gives the best performance (AUC_ROC of 0.940 and AUC_PR of 0.729).

### 3.2 Effect of ensemble learning

We trained the convolutional neural network twenty times and the ten top-performing neural networks (Models in Table 4) were chosen from performance assessment on the Val597 dataset. Selecting more models did not contribute to an increase in accuracy (data not shown). Then we averaged predictions of the 10 top-performing models to produce the ensemble for Val597, CASP10 and Val200 datasets (last row of Table 4).

**Table 4.** Effect of ensemble learning on three validation datasets*.

| Models | Val597 |  | CASP10 |  | Val200 |  |
| --- | --- | --- | --- | --- | --- | --- |
|  | AUC ROC | AUC PR | AUC ROC | AUC PR | AUC ROC | AUC PR |
| 1 | <u>0.940</u> | 0.729 | 0.892 | 0.571 | <u>0.942</u> | 0.799 |
| 2 | 0.936 | 0.711 | <u>0.897</u> | 0.555 | <u>0.942</u> | 0.798 |
| 3 | 0.935 | 0.729 | 0.889 | 0.542 | 0.934 | 0.790 |
| 4 | 0.934 | <u>0.735</u> | 0.894 | <u>0.589</u> | 0.936 | <u>0.810</u> |
| 5 | 0.931 | 0.723 | 0.888 | 0.531 | 0.935 | 0.798 |
| 6 | 0.930 | 0.719 | 0.889 | 0.557 | 0.931 | 0.792 |
| 7 | 0.930 | 0.712 | 0.894 | 0.575 | 0.937 | 0.805 |
| 8 | 0.929 | 0.719 | 0.889 | 0.547 | 0.932 | 0.803 |
| 9 | 0.929 | 0.700 | 0.882 | 0.513 | 0.931 | 0.772 |
| 10 | 0.927 | 0.716 | 0.882 | 0.502 | 0.925 | 0.794 |
| Ensemble | **0.948** | **0.776** | **0.917** | **0.606** | **0.950** | **0.841** |

\* Ten models are numbered according to the order of AUC\_ROC values for the Val597 dataset.

As shown in Table 4, the ensemble leads to a clear significant improvement over any single component model across all three datasets. For example, AUC_ROC values of the ensemble are improved by 0.8%, 2.0% and 0.8% over the highest-performing component models, for three validation sets of Val597, CASP10 and Val200, respectively. Improvement on AUC_PR is much stronger than on AUC_ROC, with 4.1%, 1.7% and 3.1% for the three validation sets, respectively.

The careful observation of AUC_ROC or AUC_PR values in Table 4 reveals that the ten models perform differently in three validation sets. For example, Models 1, 2, and 3 are the three top-performing models for the Val597 set, but Models 2, 4 and 7 are the top-three models for CASP10 dataset. This shows the variation in model ranking on different validation sets, which emphasizes the necessity of model ensembling.

### 3.3 Effect of the smoothing

We smoothed the outputs of the ensemble by averaging the predictions of the sliding window centered on the interested residue. We tested window size (w) ranging from 3 to 23 for three validation sets (Table 5).

**Table 5.** Effect of the smoothing on three validation datasets.

| Window size | Val597 |  | CASP10 |  | Val200 |  |
| --- | --- | --- | --- | --- | --- | --- |
|  | AUC ROC | AUC PR | AUC ROC | AUC PR | AUC ROC | AUC PR |
| Ensemble | 0.948 | 0.776 | 0.917 | 0.606 | 0.950 | 0.841 |
| 3 | 0.949 | **0.777** | 0.919 | **0.607** | 0.951 | 0.844 |
| 5 | 0.950 | **0.777** | 0.921 | **0.607** | 0.952 | 0.848 |
| 7 | 0.950 | 0.776 | **0.922** | 0.606 | 0.954 | 0.852 |
| 9 | **0.951** | 0.775 | **0.922** | 0.604 | 0.955 | 0.855 |
| 11 | **0.951** | 0.770 | 0.921 | 0.600 | 0.957 | 0.859 |
| 13 | 0.950 | 0.765 | 0.920 | 0.596 | 0.958 | 0.862 |
| 15 | 0.950 | 0.759 | 0.918 | 0.589 | 0.959 | 0.864 |
| 17 | 0.948 | 0.752 | 0.915 | 0.582 | 0.960 | 0.865 |
| 19 | 0.947 | 0.743 | 0.912 | 0.575 | **0.961** | **0.866** |
| 21 | 0.946 | 0.734 | 0.909 | 0.567 | **0.961** | 0.865 |
| 23 | 0.944 | 0.725 | 0.906 | 0.559 | 0.960 | 0.864 |

AUC_ROC value increases with the window starting from size 3, reaches the best at a certain size and starts to decrease as the window size continues to increase in all three validation sets. However, the window size effect on AUC_PR is different from that of AUC_ROC values. When the window size increases from 3 to 23, AUC_PR values continue to decrease in Val597 and CASP10 datasets whereas it increases up to window size 19 in Val200 dataset.

According to Table 5, the best window size for the Val597 dataset is 9, at which AUC_ROC value (0.951) of the final model is slightly higher than that (0.948) of the ensemble, and AUC_PR (0.775) is slightly lower than that (0.776) of the ensemble. CASP10 dataset has the best window size of 7, at which AUC_ROC value (0.922) of the final model is higher than that (0.917) of the ensemble and AUC_PR (0.606) equals the ensemble. For the Val200 dataset, the best window size is 19, at which two measures (AUC_ROC 0.961, AUC_PR 0.866) of the final model are much higher than those (0.950, 0.841) of the ensemble.

The best window sizes are 9, 7, and 19 for Val597, CASP10 and Val200 dataset, respectively, indicating that the different datasets are indeed different. It is thought that the best window size is proportional to the percentage of disordered residues in the dataset. Based on our experiment, the higher the fraction of disordered residues, the longer the best window size for the smoothing is. Indeed, CASP10 dataset with the low fraction (5.92%) of disordered residues has the shortest window size of 7. In contrast, in the Val200 dataset with a higher fraction (11.04%) of disordered residues, the best performance is achieved with the longest window size (w=19). In the Val597 dataset with 7.95% of disordered residues, the best window size for the smoothing is 9, which is between 7 and 19. The effect of the smoothing is more remarkable for the Val200 dataset and weaker for the Val597 dataset.

In addition, performance in three validation sets also seems to be related to the percentage of disordered residues in the dataset. From our experiment, the higher the fraction of disordered residues, the better the performance of the validation set is. In their best window sizes (7, 9, 19), the AUC_ROC (AUC_PR) values of CASP10, Val597 and Val200 dataset are 0.922 (0.606), 0.951 (0.775) and 0.961 (0.866), respectively, which are proportional to the percentage of disordered residues in the datasets which is 5.92%, 7.95% and 11.04%, respectively.

The final prediction obtained by smoothing with a window size of 9 is called PredIDR_short, whereas the final prediction obtained by smoothing with a window size of 19 is called PredIDR_long.

### 3.4 Comparison to methods participating in CAID1

Before CAID2, we evaluated performance of PredIDR_short and PredIDR_long on CAID1-DisProt-PDB dataset provided by CAID1 organizers and compared them with methods participating in CAID1 (Table 6). The predictions of other methods and interactive figures with the curves for all other methods of CAID1 are available at URL: https://caid.idpcentral.org/challenge/results.

**Table 6.** Comparison to methods participating in CAID1 for CAID1-DisProt-PDB dataset*.

| Rank | Methods | AUC ROC | AUC PR |
| --- | --- | --- | --- |
| **1** | **PredIDR-long** | **0.915** | **0.863** |
| **2** | **PredIDR-short** | **0.908** | **0.855** |
| 3 | SPOT-Disorder-Single | 0.896 | 0.835 |
| 4 | RawMSA | 0.894 | 0.812 |
| 5 | DISOPRED-3.1 | 0.875 | 0.801 |
| 6 | IsUnstruct | 0.868 | 0.774 |
| 7 | AUCpred-np | 0.864 | 0.798 |
| 8 | IUPred2A-long | 0.862 | 0.784 |
| 9 | IUPred2A-short | 0.858 | 0.758 |
| 10 | DisoMine | 0.858 | 0.711 |
| 11 | PyHCA | 0.822 | 0.707 |
| 12 | DisPredict-2 | 0.667 | 0.478 |
| 13 | DFLpred | 0.38 | 0.269 |

\* Only the methods provided predictions over all 646 sequences are shown.

31 IDR prediction methods participated in CAID1, but only 11 methods provided outputs over all 646 sequences. PredIDR_long and PredIDR_short outperform 11 IDR prediction methods in both measures.

### 3.5 Comparison to methods participating in CAID2

We took part in the second round of Critical Assessment of protein Intrinsic Disorder (CAID2) in 2022 with two programs: PredIDR_short and PredIDR_long. Table 7 shows a comparison of our disorder prediction methods with other methods participating in CAID2 for CAID2-Disorder-PDB dataset. Interactive figures with the curves for all methods of CAID2 are also available in the CAID Website (https://caid.idpcentral.org/challenge/results). According to CAID2 assessors^[5]^, some methods did not output a prediction under the given conditions because of unexpected crashing or very slow speed. So, only 28 of all 40 IDR prediction methods participated in CAID2 provided an output for all 348 sequences.

**Table 7.** Comparison to methods participating in CAID2 for CAID2-Disorder-PDB dataset*.

| Rank | Methods | AUC ROC | AUC PR |
| --- | --- | --- | --- |
| **1** | **PredIDR-long** | **0.933** | **0.87** |
| 2 | SPOT-Disorder | 0.931 | 0.889 |
| 3 | SETH-0 | 0.93 | 0.894 |
| **4** | **PredIDR-short** | **0.927** | **0.859** |
| 5 | Metapredict | 0.923 | 0.878 |
| 6 | DeepIDP-2L | 0.922 | 0.858 |
| 7 | SPOT-Disorder-Single | 0.917 | 0.87 |
| 8 | SETH-1 | 0.911 | 0.854 |
| 9 | rawMSA | 0.91 | 0.838 |
| 10 | AIUPred | 0.903 | 0.855 |
| 11 | ESpritz-D | 0.898 | 0.81 |
| 12 | pyHCA | 0.898 | 0.825 |
| 13 | VSL2 | 0.887 | 0.813 |
| 14 | Disomine | 0.886 | 0.778 |
| 15 | IsUnstruct | 0.885 | 0.799 |
| 16 | IUPred3 | 0.885 | 0.825 |
| 17 | fIDPnn | 0.882 | 0.804 |
| 18 | DISOPRED3-diso | 0.879 | 0.809 |
| 19 | ESpritz-X | 0.878 | 0.793 |
| 20 | AUCpred-no-profile | 0.876 | 0.815 |
| 21 | Dispredict3 | 0.873 | 0.794 |
| 22 | MobiDB-lite | 0.868 | 0.801 |
| 23 | ESpritz-N | 0.859 | 0.766 |
| 24 | RONN | 0.857 | 0.764 |
| 25 | DisEMBL-dis465 | 0.792 | 0.653 |
| 26 | s2D | 0.771 | 0.519 |
| 27 | DisEMBL-disHL | 0.702 | 0.5 |
| 28 | APOD | 0.537 | 0.377 |

\* Only the methods provided predictions over all 348 sequences are shown.

Among those methods, PredIDR_long ranked first and fourth and PredIDR_short fourth and sixth when considering AUC_ROC and AUC_PR, respectively.

In CAID1-DisProt-PDB and CAID2-Disorder-PDB datasets, with a disorder percentage of 16.22% and 12.92%, the performance of PredIDR_long is better than PredIDR_short (Table 6 and 7), confirming that PredIDR_long performs better than PredIDR_short on a dataset with a higher percentage of disordered residues. Similarly, PredIDR_long is better on the Val200 dataset which contains more disordered residues compared with the other datasets evaluated in Table 5.

According to CAID2 challenge^[5]^, disordered regions of the CAID2-Disorder-PDB dataset come from all types of experimental evidence reported in DisProt, including circular dichroism and X-ray missing residues. We tested methods’ ability to detect X-ray missing residues and the other sources of disorder separately. To do this, we built the CAID2-Xray-PDB and CAID2-NoX-PDB datasets (see Section 2.2).

Table 8 shows comparison of our methods with other CAID2 methods for the CAID2-Xray-PDB dataset. Among 29 methods which provided predictions over all 138 protein sequences, PredIDR_short and PredIDR_long ranked first and second for both measures.

**Table 8.**
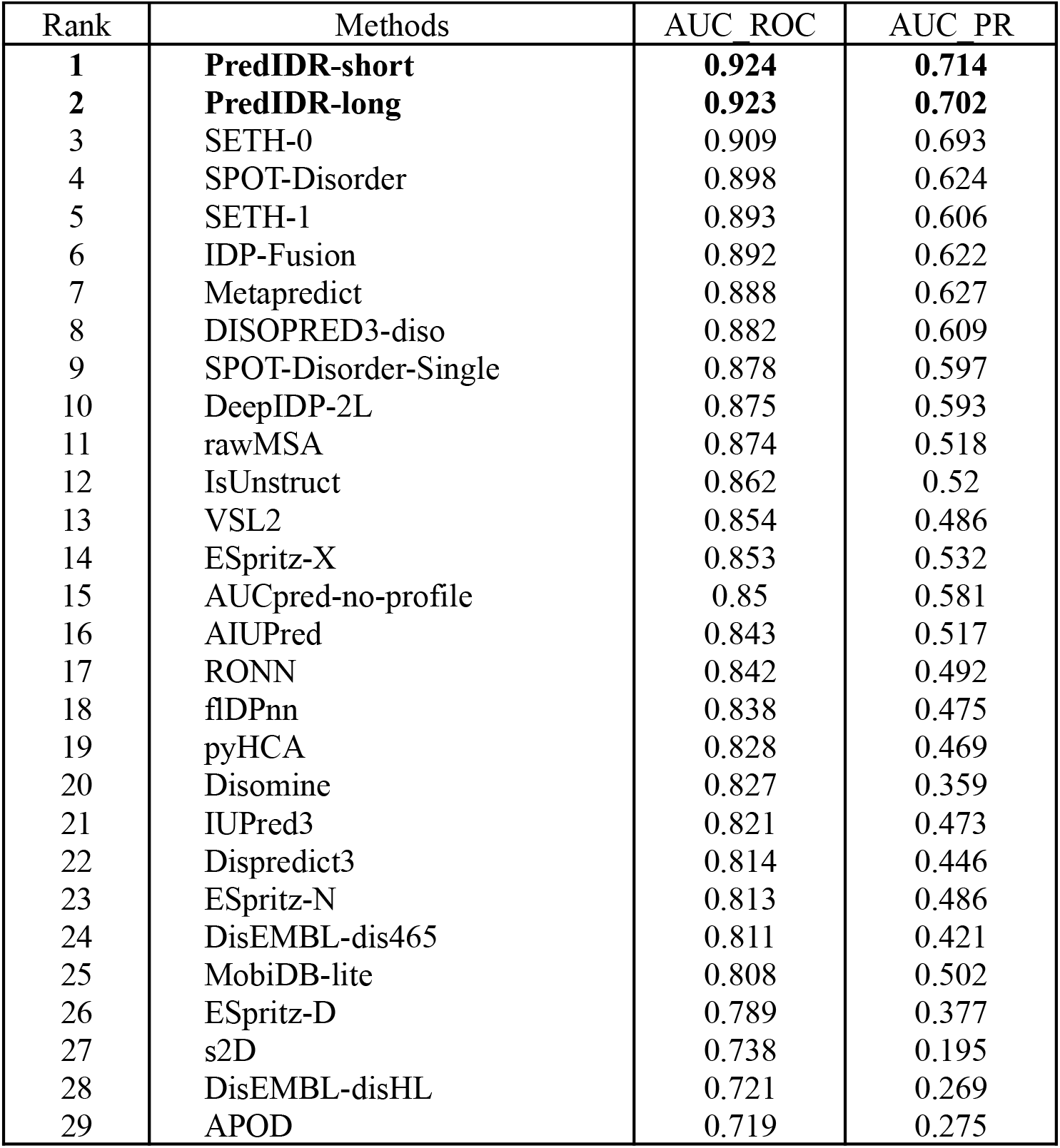
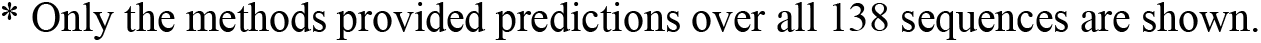
Comparison to methods participating in CAID2 for CAID2-Xray-PDB dataset*.

| Rank | Methods | AUC_ROC | AUC_PR |
| --- | --- | --- | --- |
| **1** | **PredIDR-short** | **0.924** | **0.714** |
| **2** | **PredIDR-long** | **0.923** | **0.702** |
| 3 | SETH-0 | 0.909 | 0.693 |
| 4 | SPOT-Disorder | 0.898 | 0.624 |
| 5 | SETH-1 | 0.893 | 0.606 |
| 6 | IDP-Fusion | 0.892 | 0.622 |
| 7 | Metapredict | 0.888 | 0.627 |
| 8 | DISOPRED3-diso | 0.882 | 0.609 |
| 9 | SPOT-Disorder-Single | 0.878 | 0.597 |
| 10 | DeepIDP-2L | 0.875 | 0.593 |
| 11 | rawMSA | 0.874 | 0.518 |
| 12 | IsUnstruct | 0.862 | 0.52 |
| 13 | VSL2 | 0.854 | 0.486 |
| 14 | ESpritz-X | 0.853 | 0.532 |
| 15 | AUCpred-no-profile | 0.85 | 0.581 |
| 16 | AIUPred | 0.843 | 0.517 |
| 17 | RONN | 0.842 | 0.492 |
| 18 | fIDPnn | 0.838 | 0.475 |
| 19 | pyHCA | 0.828 | 0.469 |
| 20 | Disomine | 0.827 | 0.359 |
| 21 | IUPred3 | 0.821 | 0.473 |
| 22 | Dispredict3 | 0.814 | 0.446 |
| 23 | ESpritz-N | 0.813 | 0.486 |
| 24 | DisEMBL-dis465 | 0.811 | 0.421 |
| 25 | MobiDB-lite | 0.808 | 0.502 |
| 26 | ESpritz-D | 0.789 | 0.377 |
| 27 | s2D | 0.738 | 0.195 |
| 28 | DisEMBL-disHL | 0.721 | 0.269 |
| 29 | APOD | 0.719 | 0.275 |

\* Only the methods provided predictions over all 138 sequences are shown.

However, for the CAID2-NoX-PDB dataset, PredIDR_long ranked seventh and eleventh in terms of AUC_ROC and AUC_PR, considering the 28 methods which provided predictions for all 188 protein sequences (Table 9). PredIDR_short ranked twelfth and sixteenth.

**Table 9.** Comparison to prediction methods participating in CAID2 for CAID2-NoX-PDB dataset*.

| Rank | Methods | AUC ROC | AUC PR |
| --- | --- | --- | --- |
| 1 | SPOT-Disorder | 0.933 | 0.936 |
| 2 | SETH-0 | 0.93 | 0.938 |
| 3 | DeepIDP-2L | 0.923 | 0.917 |
| 4 | Metapredict | 0.923 | 0.926 |
| 5 | SPOT-Disorder-Single | 0.92 | 0.927 |
| 6 | rawMSA | 0.915 | 0.916 |
| 7 | **PredIDR-long** | **0.911** | **0.895** |
| 8 | AIUPred | 0.911 | 0.92 |
| 9 | SETH-1 | 0.911 | 0.913 |
| 10 | ESpritz-D | 0.91 | 0.886 |
| 11 | pyHCA | 0.909 | 0.906 |
| 12 | **PredIDR-short** | **0.902** | **0.884** |
| 13 | Disomine | 0.896 | 0.879 |
| 14 | fIDPnn | 0.896 | 0.895 |
| 15 | IUPred3 | 0.894 | 0.903 |
| 16 | VSL2 | 0.89 | 0.893 |
| 17 | Dispredict3 | 0.889 | 0.89 |
| 18 | IsUnstruct | 0.888 | 0.884 |
| 19 | ESpritz-X | 0.877 | 0.875 |
| 20 | AUCpred-no-profile | 0.872 | 0.884 |
| 21 | MobiDB-lite | 0.87 | 0.878 |
| 22 | ESpritz-N | 0.858 | 0.858 |
| 23 | RONN | 0.857 | 0.857 |
| 24 | DISOPRED3-diso | 0.856 | 0.86 |
| 25 | DisEMBL-dis465 | 0.786 | 0.775 |
| 26 | s2D | 0.77 | 0.701 |
| 27 | DisEMBL-disHL | 0.695 | 0.661 |
| 28 | APOD | 0.502 | 0.518 |

\* Only the methods provided predictions over all 188 sequences are shown.

We drowned three important conclusions from the comparison of Table 8 and 9. First, most of CAID2 methods (except APOD, PredIDR, DisEMBL-disHL, DisEMBL-dis465, DISOPRED3-diso), perform better, both in terms of AUC_ROC and AUC_PR values in CAID2-NoX-PDB than CAID2-Xray-PDB dataset. For instance, the AUC_ROC and AUC_PR values of SPOT-Disorder are 0.933 and 0.936, in CAID2-NoX-PDB, 0.898 and 0.624 in CAID2-Xray-PDB dataset. The performance of the prediction methods seems to be related to the fraction of the disordered residues in the two testing sets, 4.10% in CAID2-Xray-PDB and 19.58% in CAID2-NoX-PDB as confirmed by the results of Table 5.

Second, the ranking of PredIDR_short and PredIDR_long is different in the two testing sets. On CAID2-Xray-PDB with a low fraction (4.10%) of disordered residues, PredIDR_short performs better, whereas PredIDR_long performs better on the CAID2-NoX-PDB dataset containing a higher fraction (19.58%) of disordered residues. This result is well explained in Table 5, where PredIDR_short performs better on those datasets (Val597 and CASP10) containing a low fraction of disordered residues while PredIDR_long performs better on the Val200 dataset containing a higher fraction of disordered residues.

Last, PredIDR_short and PredIDR_long have the best performance on the CAID2-Xray-PDB, but not on CAID2-NoX-PDB, probably because they are trained only on X-ray missing residues. In order to achieve better performance on the CAID2-NoX-PDB dataset, our prediction methods should be trained by non-X-ray annotations.

In the future, we are going to make a prediction method trained on the disordered residues annotated by non-X-ray annotations as well as the X-ray missing residues, which is expected to have better performance than the current PredIDR.

## 4. Conclusions

In this paper we described a new deep learning method, PredIDR, for the prediction of protein intrinsically disordered regions. Our method is trained for the detection of X-ray missing residues in protein sequences. The convolutional neural network used in our experiment consists of two 2D convolutional layers, one 2D max-pooling layer and two fully connected layers. We examined various feature combinations, the use of an ensemble method, and smoothing on the performance of the method. Our experiments show that sequence profiles, eight-state secondary structure and 20-class solvent accessibility are useful for the prediction of disorder. An ensemble prediction generated by averaging the ten top-performing models performs better than all models alone. Finally, the smoothing technique, implemented by averaging predictions within a sliding window, provides a slight improvement over the raw ensemble prediction. It seems that the optimal window size of the sliding window depends on the fraction of disordered residues in the dataset.

The test of PredIDR on the CAID1-DisProt-PDB dataset of the CAID1 experiment shows that PredIDR_long and PredIDR_short remarkably outperform 11 CAID1 methods.

Finally, our method, PredIDR, took part in the second round of the Critical Assessment of protein Intrinsic Disorder (CAID2) in 2022 performing as or better than the other state-of-the-art prediction methods^[5]^.

PredIDR can be freely used through the CAID Prediction Portal^[1]^ available at URL: https://caid.idpcentral.org/portal.

## Acknowledgements

This work was supported by ELIXIR, the research infrastructure for life science data. DP is supported by the European Union through NextGenerationEU, PNRR project National Center for Gene Therapy and Drugs based on RNA Technology (CN00000041). Italian Ministry of Education and Research through the NextGenerationEU fund PRIN 2022 project: PLANS (2022W93FTW). Funding for open access charge: University of Padova.

